# Patient-Derived Liver Cancer Organoids Reflect Tumor Biology and Their Growth Phenotype Correlates with Clinical Outcomes

**DOI:** 10.64898/2026.08.02.742347

**Authors:** Yeon Su Kim, Yoon-Ha Go, Hyung Sun Kim, Jaehwi Seo, Dong Oh Kim, Dong-Youn Hwang, Jongman Yoo, Wookyeom Yang, Jin Hong Lim

## Abstract

Liver cancer remains a major global health burden with high mortality and limited treatment response prediction tools. Patient-derived cancer organoids have emerged as promising preclinical models that recapitulate tumor heterogeneity; however, the biological significance of morphological diversity within established organoids remains poorly characterized in hepatocellular carcinoma (HCC). In this exploratory study, we investigated whether distinct organoid growth phenotypes reflect underlying tumor biology and correlate with clinical outcomes. We established liver cancer organoids from resected tumor tissues of 27 patients and analyzed their clinical, histological, and genomic characteristics. Organoids were classified morphologically into cystic and solid types. Whole exome sequencing (WES) was conducted on six matched tumor–organoid pairs to assess genomic fidelity. Associations between organoid establishment, growth characteristics, and clinical parameters were statistically evaluated. Progression-free survival (PFS) was analyzed using the Kaplan–Meier method and univariate Cox proportional hazards regression. Organoids were successfully established in 13 of 27 cases (48.1%). Solid-type organoids were significantly associated with shorter PFS compared to cystic types (HR = 13.91; p = 0.0039, log-rank test). Organoid establishment was more frequent in older patients (>70 years), those with HBV infection, and tumors with positive β-catenin expression. WES analysis demonstrated high concordance in somatic mutation profiles and variant allele frequency distributions between tissues and corresponding organoids. In univariate Cox regression, organoid growth pattern (solid vs. cystic) showed a significant association with PFS within this exploratory cohort (p = 0.0207). Patient-derived liver cancer organoids preserved the genomic and histopathological features of the original tumors. Notably, solid morphology was associated with shorter PFS, suggesting that organoid growth phenotype may serve as a supplementary indicator of tumor biological behavior in HCC. Given the modest cohort size and the absence of multivariate analysis, these preliminary findings should be interpreted with caution and warrant further large-scale validation.

## Introduction

Hepatocellular carcinoma (HCC) is one of the most common types of primary liver cancer and is considered a major global health concern [1, 2]. Each year, more than 905,677 new cases and over 830,180 deaths are reported, making HCC the third leading cause of cancer-related mortality worldwide [3]. Despite the implementation of screening programs and the development of molecular targeted therapies and immune checkpoint inhibitors, the overall 5-year survival rate remains below 20%. This poor prognosis is largely due to the asymptomatic presentation of early-stage disease, which often leads to late-stage diagnosis and limited treatment opportunities [4].

The most prominent etiologic drivers of HCC are chronic infections with hepatitis B virus (HBV) and hepatitis C virus (HCV), which together account for the majority of global HCC cases. These infections contribute to hepatocarcinogenesis through chronic inflammation, fibrosis, and eventual cirrhosis [2, 5]. Other major contributors include alcohol-related liver disease and metabolic dysfunction–associated liver conditions, such as non-alcoholic fatty liver disease (NAFLD) and non-alcoholic steatohepatitis (NASH), which are increasingly recognized as important risk factors in both Western and Asian populations [6].

Beyond these environmental and metabolic risk factors, HCC is characterized by significant molecular heterogeneity, reflective of its diverse underlying etiologies. At the genomic, epigenomic, and transcriptomic levels, HCC tumors display substantial variability [7, 8]. Frequently mutated genes such as TERT, CTNNB1, and TP53, along with amplification of oncogenes like FGF19 and MET, define distinct molecular subtypes with variable biological behavior and therapeutic response [8].

However, despite advances in molecular profiling, current clinical management of HCC still relies heavily on staging systems such as TNM and BCLC. These traditional frameworks often fail to adequately capture the underlying biological complexity of the disease, which may partly explain the limited predictive power of existing prognostic models [9, 10].

To address these limitations, various preclinical models have been used to study the pathogenesis and therapeutic response of HCC. While two-dimensional cell lines and animal models are widely used, they lack the ability to fully recapitulate the tumor microenvironment and heterogeneity [11, 12]. In contrast, patient-derived xenograft (PDX) models retain the histological and molecular characteristics of the original tumor and provide a more clinically relevant platform [13]. Nevertheless, PDX models are limited by long establishment times (typically 4–6 months), high costs, and relatively low engraftment rates for HCC (30–40%) [14]. Engraftment success is also influenced by tumor grade, vascular invasion, and degree of differentiation [13].

Moreover, the widespread use of PDX models raises ethical concerns regarding animal welfare, particularly due to the need for immunodeficient mice and the high number of animals required for successful engraftment and serial passaging [15]. These challenges underscore the need for alternative platforms that are both biologically relevant and ethically sustainable.

In this context, patient-derived tumor organoids have emerged as a promising three-dimensional culture system that may overcome the limitations of conventional models [11, 16, 17]. Organoids derived from tumor tissues can preserve key features of the original cancer, including histoarchitecture, genomic alterations, and functional properties, although their success depends on tissue quality and culture conditions. They are amenable to cryopreservation and long-term passaging and have been widely applied in drug screening, gene editing, and personalized therapeutic strategies [18].

Interestingly, emerging evidence suggests that the phenotypic characteristics of established preclinical models, rather than mere technical success, may reflect intrinsic tumor biology and aggressiveness. For example, tumors with high histologic grade, vascular invasion, and elevated expression of markers like GPC3 or Ki-67 are more likely to successfully engraft in PDX models [13, 14, 19]. Similar correlations have been observed in organoid models of pancreatic [20], colorectal [21], and breast cancers [22], where successful culture was often linked to poor clinical outcomes. These findings collectively suggest that successful model establishment may be linked to more aggressive tumor biology and poorer clinical outcomes. Moreover, recent evidence in other solid tumors suggests that post-establishment morphological or functional diversity within organoid cultures may additionally encode critical prognostic information. However, the biological significance of heterogeneous growth patterns in patient-derived HCC organoids remains poorly understood.

Given the growing clinical interest in organoid-based platforms, it is valuable to explore whether distinct morphological phenotypes arising during organoid culture reflect specific tumor characteristics. Elucidating the relationship between these early-stage cultural phenotypes and original tumor aggressiveness could expand our understanding of HCC biological heterogeneity.

In this exploratory study, we evaluated patient-derived HCC organoids to examine whether their ex vivo growth characteristics align with the genomic and histopathological features of parental tumors. Concurrently, we investigated whether structural variations in organoid growth— specifically cystic versus solid phenotypes—correlate with clinicopathological parameters and patient outcomes. Our preliminary observations suggest that a solid morphological subtype may be associated with more aggressive biological behavior and potentially unfavorable progression-free survival. While limited by a modest cohort size, these findings offer initial insights into the potential relevance of organoid morphology as a supplementary indicator of tumor behavior, warranting further large-scale validation.

## Materials and Methods

### Patient enrollment and generation of liver cancer organoids

A total of 27 patients diagnosed with liver cancer at Gangnam Severance Hospital (Seoul, Republic of Korea) were enrolled in this study between 2023 and 2025. Prior to study initiation, institutional approval was obtained from the Institutional Review Board of Gangnam Severance Hospital (IRB approval No. 3-2023-0347), and written informed consent was acquired from all participants.

Human hepatocellular carcinoma (HCC) tissues were collected and mechanically minced, followed by enzymatic digestion using 2.5 mg/mL collagenase D (Roche) in Hanks’ Balanced Salt Solution (HBSS; Welgene) at 37°C for 1 hour. After complete dissociation, the cell suspension was filtered through a 100 μm nylon cell strainer and centrifuged at 1500 rpm for 5 minutes. The resulting pellet was washed with cold Advanced DMEM/F12 (GIBCO) and resuspended in 70% Matrigel (Corning). The Matrigel-cell mixture was plated into 48-well culture plates and incubated at 37°C with 5% CO2 for 20 minutes for matrix polymerization.

The hepatocellular carcinoma organoid (HCCO) expansion medium consisted of Advanced DMEM/F12 (GIBCO) supplemented with 10% R-spondin-1 conditioned medium (ORGANOIDSCIENCES), 1× B-27 supplement without vitamin A (GIBCO), 10 mM nicotinamide (Sigma-Aldrich), 10 mM HEPES, 1× GlutaMAX, 1× penicillin/streptomycin (Thermo Fisher Scientific), 100 ng/mL recombinant human FGF10 (ORGANOIDSCIENCES), 1× N-2 supplement (GIBCO), 1.25 mM N-acetylcysteine (Sigma-Aldrich), 50 ng/mL recombinant human EGF (ORGANOIDSCIENCES), 25 ng/mL recombinant human HGF (ORGANOIDSCIENCES), 10 μM forskolin, 10 nM human [Leu15]-gastrin I (Sigma-Aldrich), and 5 μM A83-01 (Tocris). The medium was refreshed every 2 to 3 days.

### Histology and immunohistochemistry

To evaluate the histological similarity between primary liver tumor tissues and matched patient-derived organoids, both tumor tissues and liver cancer organoids were fixed in 4% paraformaldehyde (PFA; Biosolution Co., Ltd., Seoul, Korea) for 24 hours and 30 minutes, respectively, and embedded in paraffin. Paraffin-embedded sections (4 μm thick) were deparaffinized in xylene, rehydrated through a graded ethanol series, and stained with hematoxylin and eosin (H&E) using standard protocols.

For immunohistochemistry (IHC), antigen retrieval was performed by heating the sections in sodium citrate buffer (10 mM sodium citrate with 0.05% Tween-20, pH 6.0) at 95°C for 1 hour. Endogenous peroxidase activity was quenched with 3% hydrogen peroxide (H2O2) in methanol for 10 minutes. After blocking with 1% bovine serum albumin (BSA; GenDEPOT, Barker, TX, USA) in PBS, sections were incubated overnight at 4°C with the following primary antibodies: EpCAM (1:200), HepPar-1 (1:200), AFP (1:100), CK19 (1:200), N-cadherin (1:200; BD Transduction Laboratories, Franklin Lakes, NJ, USA), and vimentin (clone V9; 1:100; Leica Biosystems). Subsequently, sections were incubated at room temperature for 30 minutes with a biotinylated secondary antibody (1:200; anti-mouse IgG or anti-rabbit IgG) and processed using the avidin-biotin complex (ABC) method with the Vectastain ABC and DAB kits (Vector Laboratories, Burlingame, CA, USA) according to the manufacturer’s instructions.

### Whole exome sequencing and bioinformatic analysis

Genomic DNA was extracted from primary liver tumor tissues and matched patient-derived organoids using standard protocols. Whole exome libraries were prepared using the Twist Human Core Exome 2.0 NGS Library Preparation Kit (Twist Bioscience, CA, USA), and sequencing was performed by Gencurix, Inc. (Seoul, Republic of Korea) on the Illumina NovaSeq 6000 platform, generating 151 bp paired-end reads.

Sequencing reads were aligned to the GRCh37 human reference genome (hg19, human_g1k_v37_decoy) using BWA-MEM. Somatic mutation calling was conducted using Mutect2 and pileup-based variant detection, resulting in variant call format (VCF) files. These VCF files were converted to Mutation Annotation Format (MAF) using the vcf2maf tool (https://github.com/mskcc/vcf2maf), incorporating Ensembl Variant Effect Predictor (Ensembl-VEP) for variant annotation.

MAF files were subsequently analyzed in R (v4.5.1) using the maftools package (v2.24.0). Multiple tumor and organoid MAFs were merged using merge_mafs() for integrated analysis. Oncoplots were generated with oncoplot() to visualize the top 30 most frequently mutated genes, with sample order manually defined to reflect tissue–organoid pairing. Mutation overlaps were assessed by extracting gene-level mutations (Hugo_Symbol) and visualized using Venn diagrams created with the VennDiagram package. To evaluate genomic similarity between tumor and organoid samples, a binary mutation matrix was constructed and Pearson correlation coefficients were calculated. The correlation heatmap was visualized using the ggcorrplot package. Tumor mutational burden (TMB) was calculated using the tmb() function in maftools, based on non-synonymous somatic mutation counts. Group-wise comparisons between tumor and organoid samples were visualized using ggplot2, and statistical significance was assessed using the Wilcoxon rank-sum test via the ggpubr package. For HLA typing, OptiType and HLA-LA were used to infer class I HLA genotypes from the WES data. The HLA types of tumor and organoid samples were compared to assess allelic concordance and sample fidelity.

### Statistical analysis

Statistical analyses were conducted to evaluate the associations between clinicopathological variables, organoid characteristics, and progression-free survival (PFS). All tests were two-sided, with p < 0.05 considered statistically significant. Values between p < 0.1 and ≥ 0.05 were interpreted as borderline significant.

Categorical variables, including age group, gender, tumor stage, histological grade, TNM stage, viral hepatitis status (HBV, HCV), recurrence, liver cirrhosis, tumor size, AFP and PIVKA levels, neoadjuvant therapy, and expression of CK19, EpCAM, p53, beta-catenin, CD34, and organoid characteristics (establishment and morphology), were analyzed using the Chi-square test to assess correlations with organoid establishment success and morphological subtype (cystic vs. solid).

Survival analysis was performed using the Kaplan–Meier method, and differences between subgroups were assessed using the log-rank test. Comparisons included TNM stage (I–II vs. III– IV), organoid establishment status (success vs. fail), and organoid growth morphology (cystic vs. solid). Median survival time, hazard ratios (HRs), and 95% confidence intervals (CIs) were calculated.

To identify prognostic factors for PFS, univariate Cox proportional hazards regression analysis was performed for the same set of clinical and experimental variables. Variables with p < 0.1 in univariate analysis were considered potentially relevant and were included in reporting. Although multivariate Cox regression was attempted, the limited sample size and missing data across covariates precluded a statistically valid model; thus, multivariate results were not included in the final interpretation.

All statistical analyses were conducted using R (version 4.5.1) with the survival, dplyr, and writexl packages. Kaplan–Meier curves and associated visualizations were generated using GraphPad Prism version 8.0 (GraphPad Software, San Diego, CA, USA).

## Results

### Establishment and morphological classification of liver cancer organoids

We attempted to generate patient-derived liver cancer organoids (PDOs) using surgically resected tumor tissues from 27 patients diagnosed with HCC. Organoid cultures were successfully established in 13 out of 27 cases, corresponding to a success rate of 48.1%. The organoids formed distinct three-dimensional structures that could be morphologically classified into two phenotypes: cystic (n = 10) and solid (n = 3) (Fig 1). Time-lapse imaging revealed that cystic organoids tended to expand with a clear central lumen, whereas solid-type organoids formed dense, compact structures without a central cavity.

**Fig 1.**
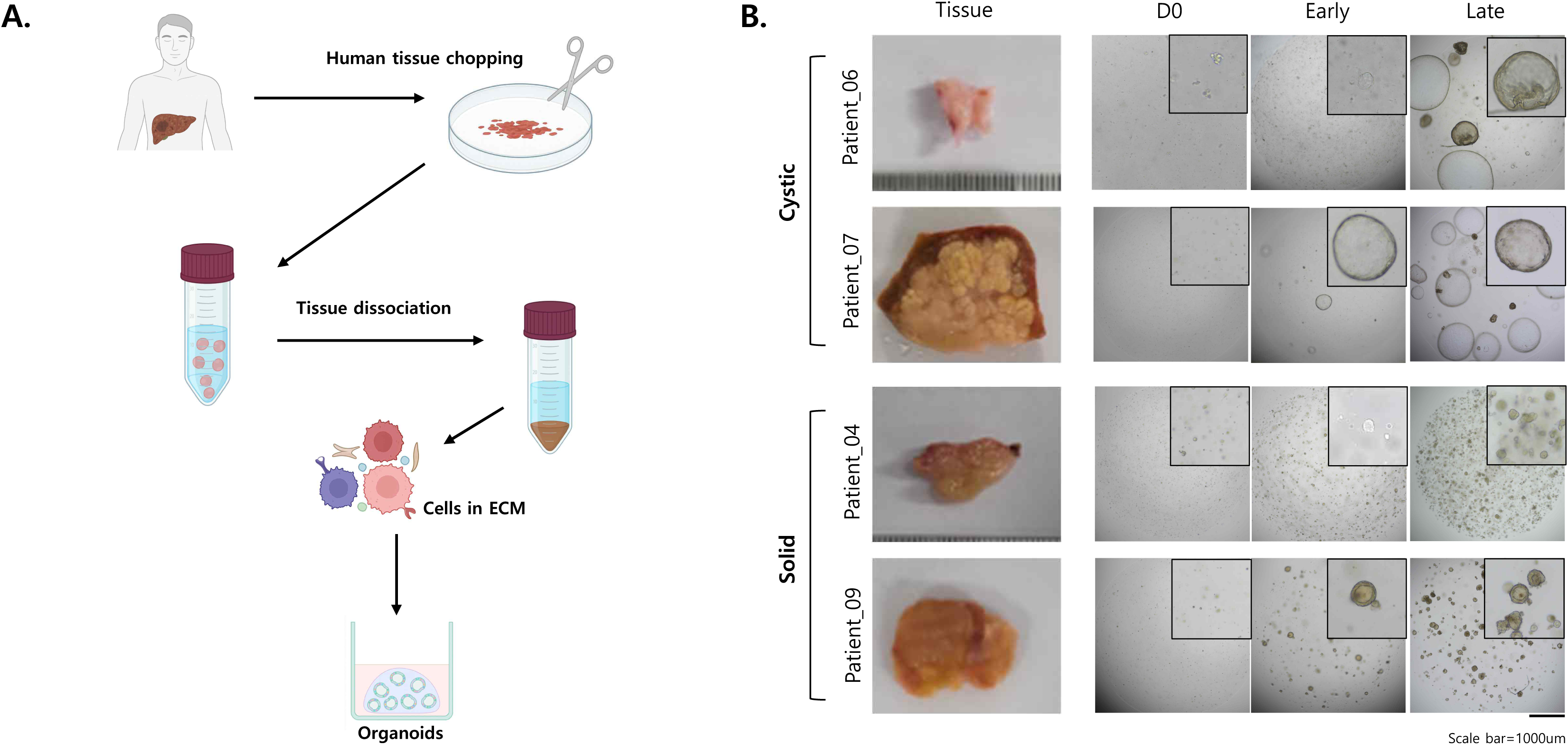
Workflow for establishment of patient-derived liver cancer organoids and representative morphology of established organoids. (A) Schematic description of the generation process of human liver cancer organoids. A total of 27 patients with HCC were enrolled, and organoids were successfully established from 13 cases (48.1%). Established organoids were morphologically classified into cystic (n = 10) and solid (n = 3) subtypes during early passages (P0–P2). The establishment timeline spans from initial tissue processing (D0) through early-passage morphological stabilization. (B) Bright-field images of liver cancer tissue and derived organoids. Labels D0, Early, and Late indicate the time points of image acquisition, which vary among organoid lines. For Patient_06 and Patient_07, images were acquired at P0D5 and P2D7; for Patient_04, at P0D7 and P2D7; and for Patient_09, at P0D7 and P2D11. Scale bar = 1000 μm.

To assess the histological fidelity of the organoids, H&E staining and immunohistochemical analysis were performed on both tumor tissues and the corresponding organoids (Fig 2). Organoids maintained the architectural integrity of the primary tumors and expressed key HCC-associated markers, including CK19, AFP, EpCAM, and HepPar-1, in patterns similar to those of the original tumor tissue, supporting the notion that liver PDOs recapitulate the phenotypic and molecular features of the parental tumors.

**Fig 2.**
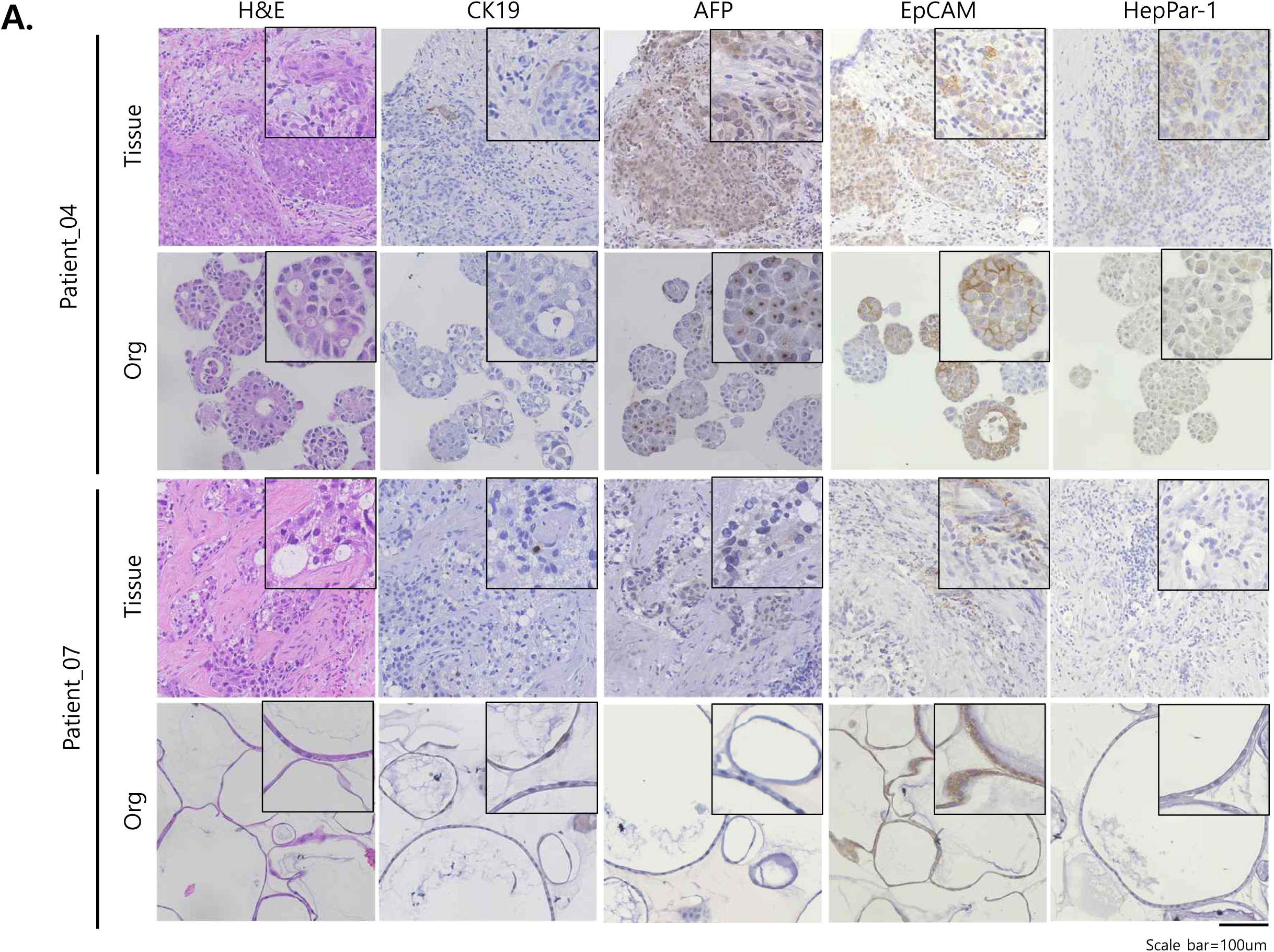
Histological and molecular similarity between patient tumors and corresponding organoids. H&E and immunohistochemistry staining of liver cancer organoids and their corresponding original tumor tissues. Organoids and tumor tissues were stained for CK19, AFP, EpCAM, and HepPar-1 to compare marker expression patterns. Scale bar = 100 μm.

### Clinicopathological factors associated with organoid establishment and morphological subtype

To investigate the clinical and pathological factors influencing the success of organoid generation, we analyzed multiple patient variables using the chi-square test (Table 1). We found that older age (≥ 70 years) was significantly associated with successful organoid formation (p = 0.0313). Furthermore, HBV-positive status was more frequently observed in the success group than in the failure group (p = 0.0217), suggesting that the viral hepatitis–associated tumor microenvironment may favor organoid formation. Importantly, tumors expressing β-catenin showed a markedly higher organoid formation rate than β-catenin–negative tumors (p = 0.0035), implicating Wnt pathway activation as a potential contributor to organoid viability and self-renewal capacity [29].

Analysis of morphological subtype revealed that solid-type organoids were significantly associated with recurrence (p = 0.0031) and tended to arise from more aggressive tumors, although no significant associations were found with TNM stage or AFP/PIVKA levels. These findings suggest that organoid morphology may reflect the biological behavior of the tumor.

### Genomic fidelity between organoids and parental tumors

To determine the extent to which organoids retained the genomic features of their parental tumors, we performed WES on six matched tissue–organoid pairs. Mutation profile comparison using oncoplots showed that the majority of frequently mutated genes were conserved between each tumor and its corresponding organoid (Fig 3A). Venn diagram analysis confirmed a high proportion of shared variants within each pair, with only a small subset of mutations being unique to either the tissue or organoid (Fig 3B).

**Fig 3.**
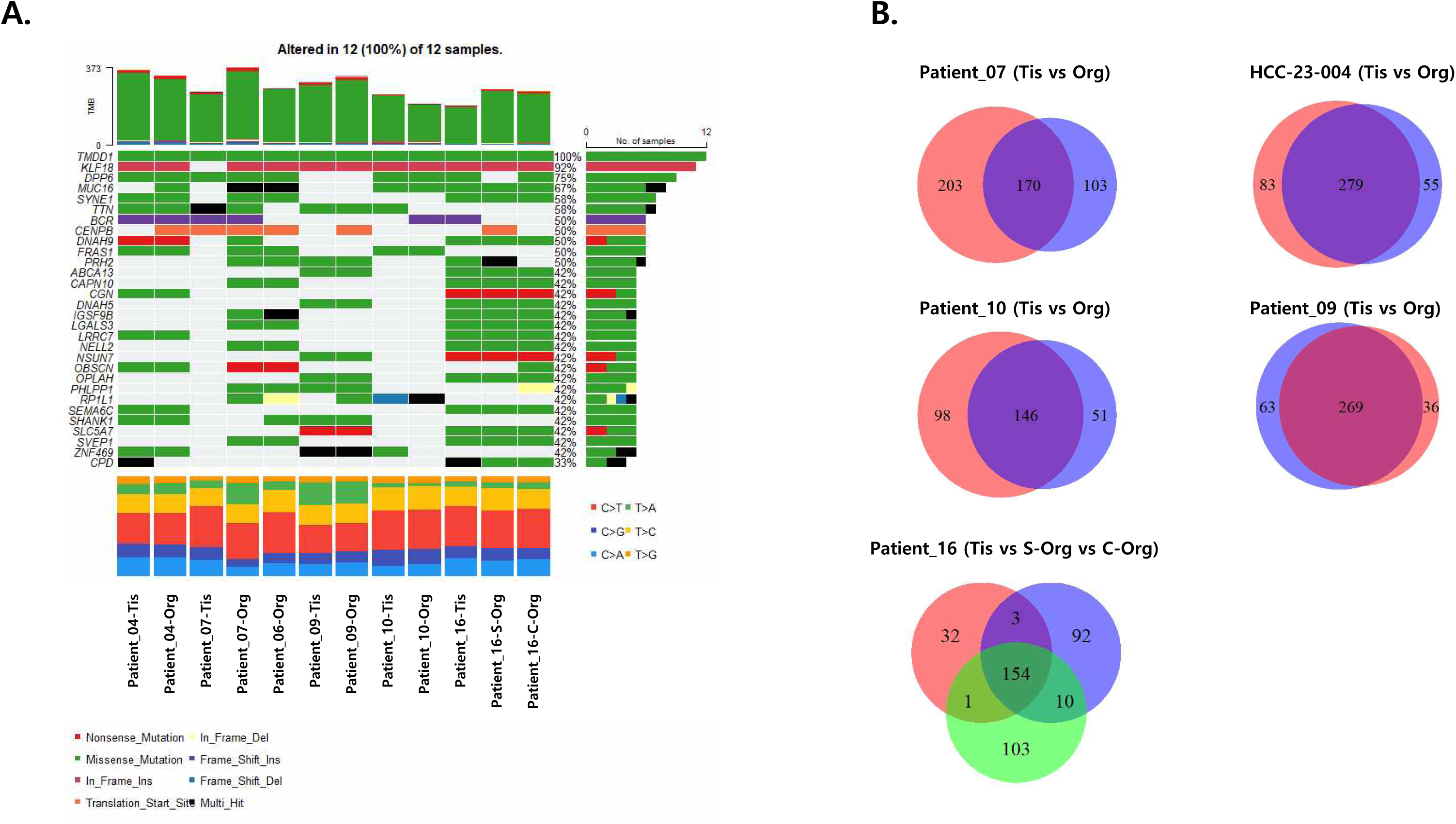
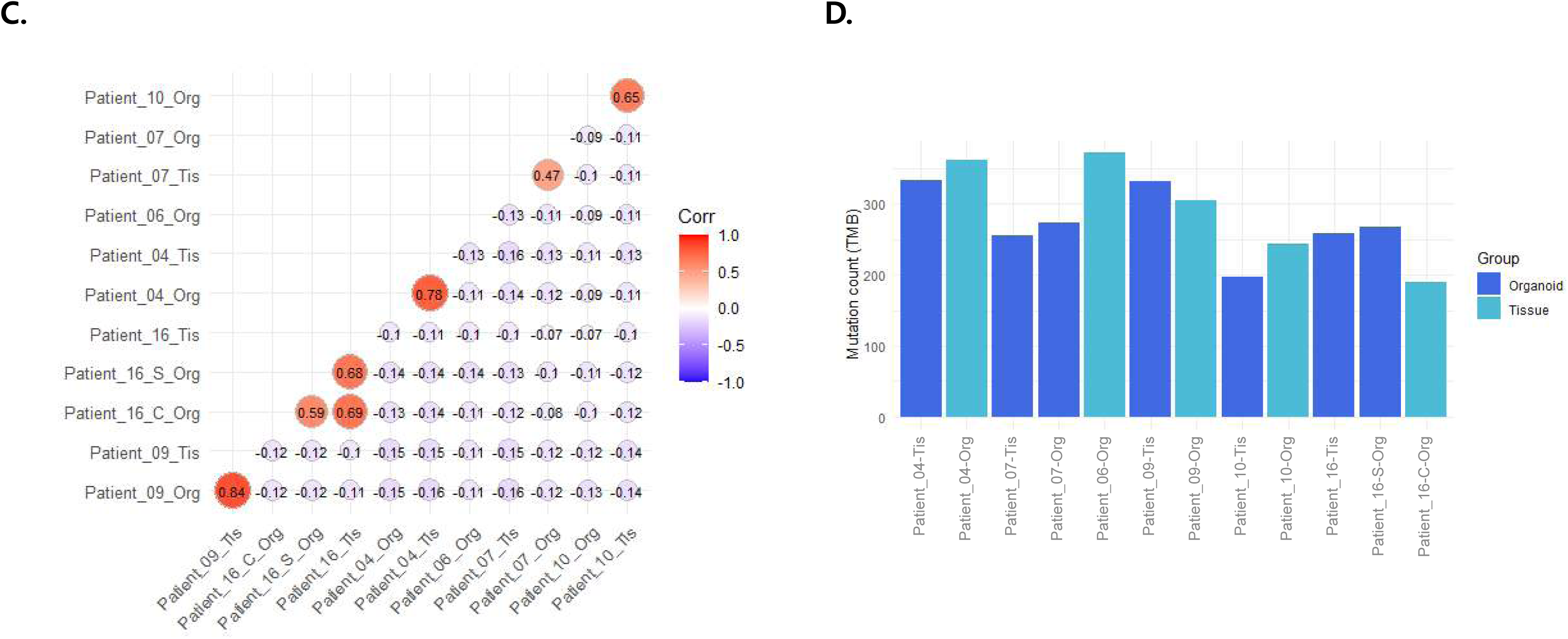
Genomic concordance between primary liver cancer tissues and matched patient-derived organoids assessed by whole exome sequencing. (A) Oncoplot showing somatic mutation profiles of primary tumor tissues and matched organoids across 12 samples. Mutation types are color-coded, and mutation frequencies across samples are summarized. (B) Venn diagrams comparing the number of shared and unique variants between tumor tissues and matched organoids for each donor. In each two-way Venn diagram, red indicates primary tumor tissue (Tis) and blue indicates matched organoid (Org). For Patient_16, a three-way comparison among tumor tissue (Tis; red), solid organoid (S-Org; green), and cystic organoid (C-Org; blue) is shown, reflecting the co-establishment of two morphologically distinct organoid lines from a single tumor. (C) Heatmap of pairwise Pearson correlation coefficients based on variant allele frequencies (VAFs) between tissue and organoid samples. (D) Bar graph comparing tumor mutational burden (TMB) between organoids and matched tumor tissues for each sample.

Furthermore, variant allele frequency (VAF) correlation analysis revealed strong concordance across all pairs, indicating that allelic distributions were preserved during the organoid derivation process (Fig 3C). Tumor mutational burden (TMB) was comparable between tissues and organoids, with no statistically significant difference observed (Fig 3D). These results collectively support that liver cancer organoids maintain the mutational landscape of the original tumors, validating their use as genetically faithful preclinical models.

Based on mutation frequency and site counts, a total of 34 genes harbored at least three independent mutations across the six HCC organoid samples (Table 2). These included DPP6, SYNE1, BCR, NELL2, and especially MUC16 (CA-125), which is associated with diverse biological functions such as membrane transport, adhesion, and cytoskeletal organization during cancer progression and metastasis [23]. While canonical HCC drivers such as TP53 and CTNNB1 were not among the most recurrently mutated genes [24], both mutations were concurrently observed in the tumor and organoid samples from Patient_09 (S1 Table), further supporting the model’s ability to preserve patient-specific genetic alterations. Of note, the VAF of TP53 (p.Y234C) increased from 0.276 in the primary tissue to 1.000 in the organoid, and that of CTNNB1 (p.K335I) increased from 0.217 to 0.475, consistent with clonal enrichment of driver mutation-bearing cells during ex vivo expansion. Collectively, these findings reinforce the genomic fidelity of patient-derived organoids and support their utility as representative preclinical models for hepatocellular carcinoma.

### HLA genotype concordance

To further confirm the genomic fidelity of organoids, we performed HLA class I (HLA-A, -B, -C) and class II (HLA-DQA1, -DQB1, -DRB1, DPA1, DPB1, DRB3, DRB4) genotyping on matched tumor and organoid samples using both OptiType and HLA-LA algorithms (Table 3). In the majority of cases, HLA types were identical between tumor and organoid, demonstrating that the organoid derivation process did not induce allelic drift or contamination. This supports the potential use of organoids in immune-related applications, including antigen presentation and T-cell interaction studies.

### Association between organoid phenotype and clinical outcome

We next evaluated whether organoid characteristics were associated with patient prognosis. Kaplan–Meier survival analysis was performed to assess PFS across different patient subgroups (Fig 4). As expected, patients with advanced TNM stage (III/IV) had significantly poorer survival than those with early-stage disease (I/II) (HR = 5.914; log-rank p = 0.0048) (Fig 4B).

**Fig 4.**
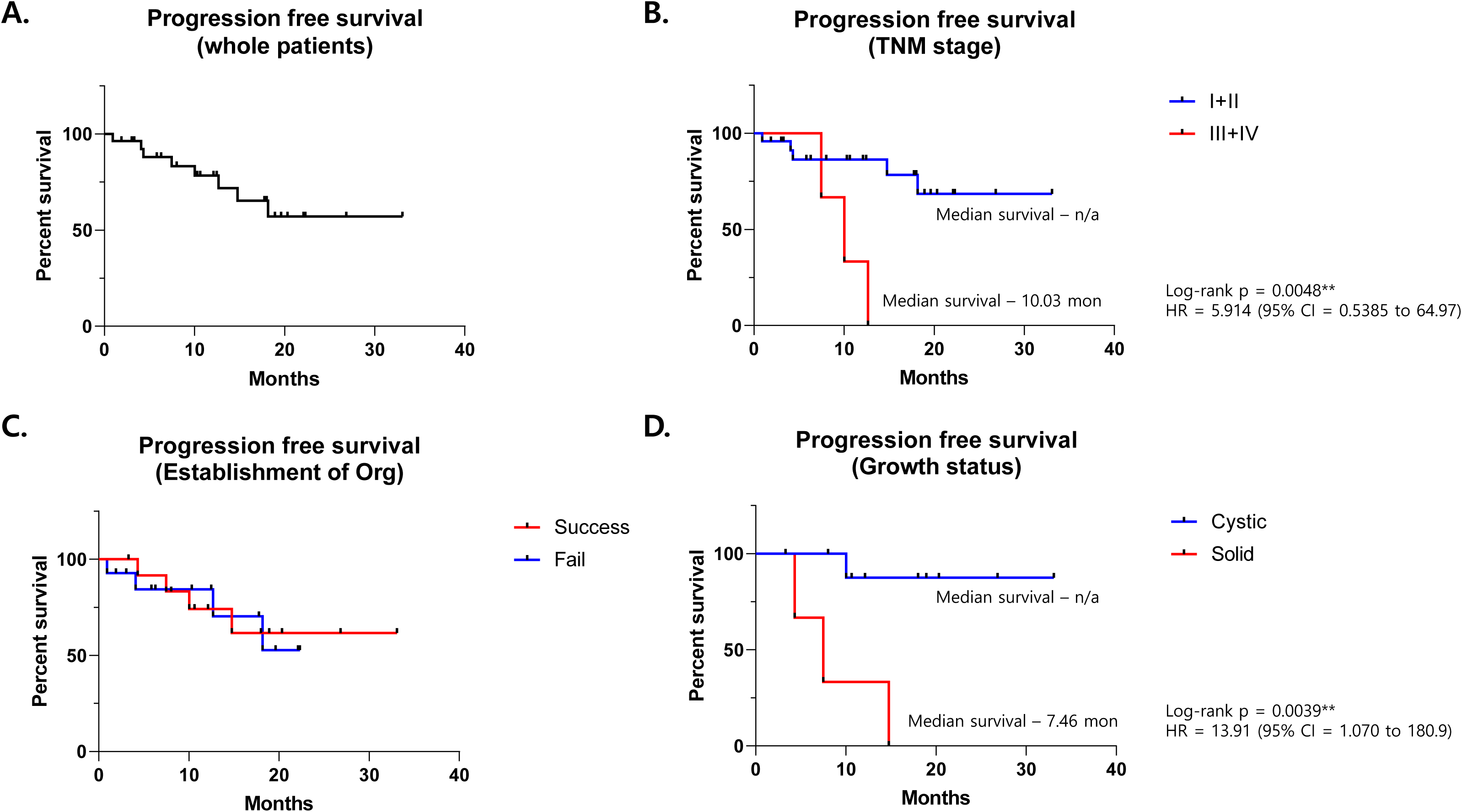
Kaplan–Meier analysis of progression-free survival (PFS) in liver cancer patients based on organoid establishment and clinicopathological characteristics. (A) PFS curve for the entire patient cohort. (B) Comparison of PFS between patients with TNM stage I/II and III/IV. Patients with advanced stage (III/IV) showed significantly poorer survival (log-rank p = 0.0048). (C) Comparison of PFS according to the success or failure of patient-derived organoid (PDO) establishment. (D) Comparison of PFS based on tumor growth status (cystic vs. solid). Patients with solid tumors exhibited significantly shorter PFS than those with cystic tumors (log-rank p = 0.0039). Statistical significance between curves was determined by the log-rank test. Hazard ratios (HR) and 95% confidence intervals (CI) are indicated.

Although organoid establishment success alone did not significantly influence survival outcomes (Fig 4C), organoid morphology showed a strong association with PFS. Patients whose tumors gave rise to solid-type organoids had markedly shorter PFS compared to those with cystic-type organoids (median PFS = 7.46 months vs. not reached; HR = 13.91; log-rank p = 0.0039) (Fig 4D). This finding suggests that solid organoid formation may be an indicator of intrinsic tumor aggressiveness.

### Prognostic value of clinical and organoid-based variables

To further explore prognostic factors, we performed univariate Cox proportional hazards regression analysis using clinical and organoid-related variables (Fig 5). Among all variables tested, organoid growth status (solid vs. cystic) was significantly associated with PFS (HR = 14.90, p = 0.0207), along with TNM stage (HR = 7.27, p = 0.0157). Tumor size (≥ 5 cm) and AFP levels (≥ 100 ng/mL) showed borderline significance (p = 0.0959 and p = 0.0983, respectively). Variables such as gender, grade, neoadjuvant therapy, and CK19/EpCAM expression were not significantly associated with clinical outcome. Due to the limited number of events and cohort size, multivariate analysis was attempted but not deemed statistically valid for inclusion (data not shown).

**Fig 5.**
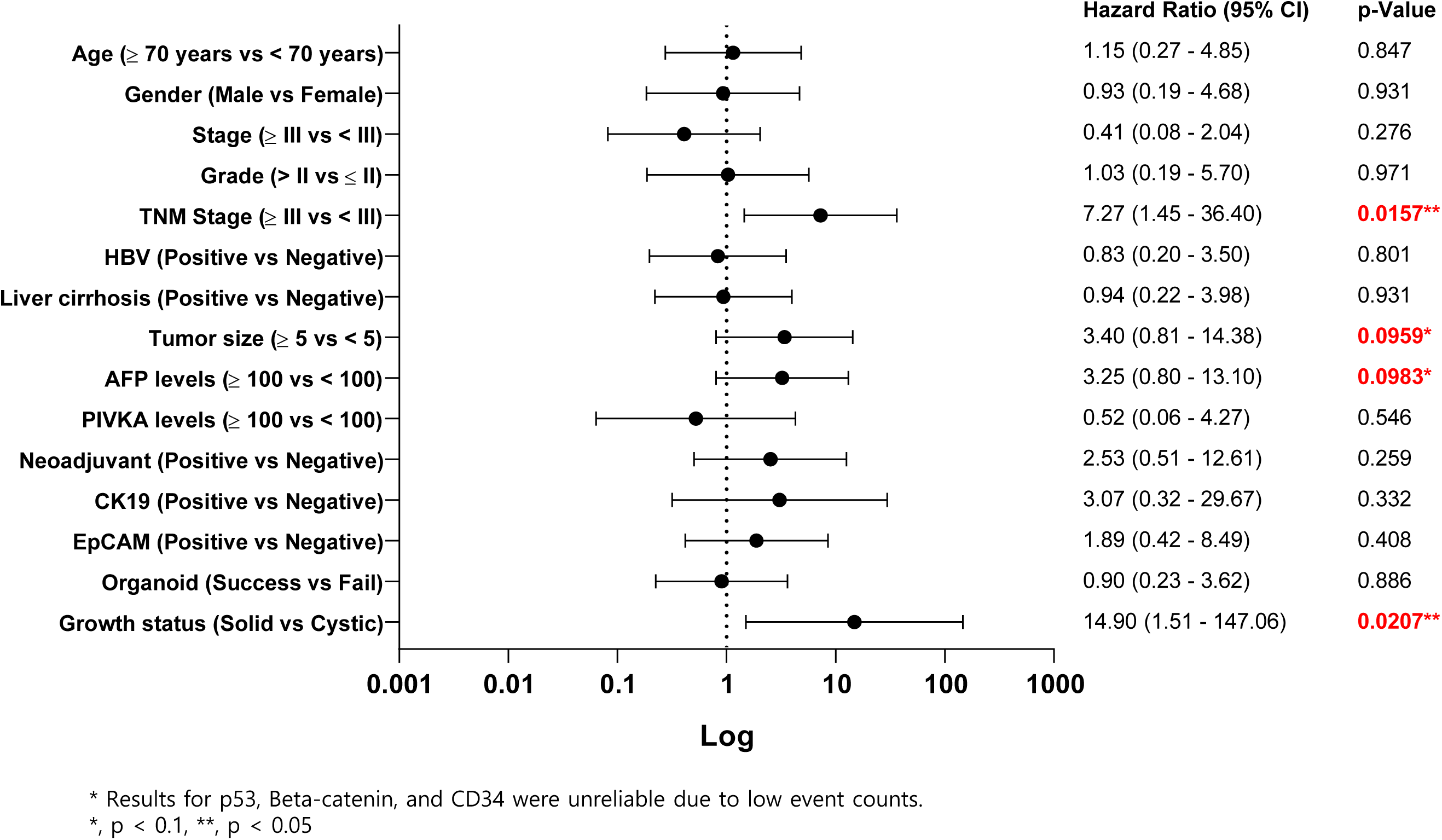
Univariate Cox regression analysis of prognostic factors in patients with liver cancer. Hazard ratios (HRs) and 95% confidence intervals (CIs) are shown on a logarithmic scale. Prognostic factors with p ≤ 0.1 were considered potentially relevant, including TNM stage, organoid growth status, tumor size, and AFP levels.

## Discussion

Hepatocellular carcinoma (HCC) remains one of the most lethal malignancies worldwide due to late diagnosis, tumor heterogeneity, and limited treatment responsiveness [1]. In efforts to better model the disease and support individualized therapeutic strategies, patient-derived tumor organoids have emerged as promising ex vivo platforms that recapitulate tumor architecture and genetic features [11]. However, the extent to which organoid characteristics correlate with clinical features and outcomes has not been thoroughly investigated in HCC. In this study, we established organoids from resected liver cancer tissues in nearly half (13/27, 48.1%) of enrolled patients and observed two distinct morphological subtypes, cystic and solid (Table 1).

Solid-type organoids were significantly associated with disease recurrence and shorter PFS, with a hazard ratio (HR) of 13.91 (log-rank p = 0.0039) (Fig 4). This result was further validated in univariate Cox regression, where solid morphology showed a significant association with poor progression-free survival (p = 0.0207) within this exploratory cohort (Fig 5). These preliminary findings suggest that organoid morphology may reflect underlying tumor features, potentially serving as a supplementary indicator of biological behavior in HCC.

While the specific mechanisms underlying cystic versus solid organoid morphology in HCC remain unclear, similar associations have been reported in colorectal cancer organoids. In those studies, solid-type organoids were linked to activation of IGF1R signaling pathways and stem-like transcriptional profiles, which are commonly associated with tumor aggressiveness [25].

Furthermore, large-scale image-based profiling has demonstrated that organoid morphology can correlate with molecular traits and drug sensitivity patterns, supporting the idea that morphological features encode meaningful biological information [26]. These observations from colorectal cancer models provide a rationale for interpreting the morphological differences observed in HCC organoids as reflective of intrinsic tumor biology. In addition, from a biophysical perspective, differences in shape may be influenced by mechanical properties such as surface tension and cell cohesion, which have been implicated in tumor invasiveness [27]. Taken together, our findings, in light of prior work in other tumor types, suggest that organoid growth phenotype, particularly solid morphology, could be leveraged as a potential indicator of aggressive features in HCC.

Interestingly, the overall organoid establishment rate in our cohort (48.1%) is comparable to reported engraftment rates in PDX models for HCC (30–40%) [19]. While PDX models retain in vivo histologic structure and offer valuable translational insights, they require immunodeficient animals, extended establishment periods, and incur high costs [14]. Furthermore, ethical concerns regarding animal welfare continue to limit their widespread use [15]. In contrast, organoid models offer a more rapid, scalable, and ethically sustainable alternative [16]. The similar establishment efficiency combined with reduced time and cost underscores the practical advantages of organoid systems for modeling liver cancer.

From a biological standpoint, we found that HBV-positive status and β-catenin expression were significantly associated with successful organoid generation. HBV-associated tumors may present with an inflammatory microenvironment or cellular composition that facilitates ex vivo viability and expansion [28], although this hypothesis warrants further functional investigation. Additionally, activation of Wnt signaling, implied by β-catenin positivity, may enhance stemness and self-renewal properties, promoting long-term growth of tumor organoids [29]. While β-catenin positivity increased organoid success rates, it was not directly linked to worse clinical outcomes in our cohort, suggesting that the role of Wnt signaling in HCC prognosis may be context-dependent.

Whole exome sequencing (WES) confirmed the genomic fidelity between tumors and their matched organoids. Across donor-matched pairs, both somatic mutation profiles and tumor mutational burden (TMB) were highly concordant. Variant allele frequency (VAF) correlations also showed strong consistency between tissues and organoids, indicating that the organoid derivation process preserved allelic distributions without introducing clonal selection biases (Fig 3C).

Crucially, to address concerns regarding the biological identity and tumor specificity of the cystic subtype, our genomic analysis provides robust evidence that these structures are genuinely tumor-derived rather than representing normal tissue overgrowth. In Patient_16, from whose primary tumor both cystic and solid organoid lines were successfully co-established, both variants demonstrated exceptional genomic fidelity to the parental tissue, retaining matching somatic mutation landscapes (Fig 3A) and exhibiting high VAF correlations (Pearson r = 0.69 for cystic, r = 0.68 for solid) (Fig 3C). This cross-validation effectively demonstrates that the cystic phenotype preserves genuine tumor-specific molecular architecture rather than representing contaminating normal hepatocytes or biliary/progenitor lines.

Notably, among the organoids analyzed, we identified frequently mutated genes across multiple cases (Table 2), including DPP6, MUC16, SYNE1, BCR, and NELL2, which are involved in membrane transport, cytoskeletal regulation, and cell adhesion. However, classical HCC driver mutations such as TP53 and CTNNB1 were only observed in one matched sample (Patient_09), limiting our ability to evaluate their broader clinical relevance within this dataset.

Moreover, HLA genotyping—including both class I and II alleles—revealed nearly complete concordance across matched samples, supporting the potential application of organoids in immunogenomic studies and personalized immunotherapy testing (Table 3). Regarding phenotypic stability across passaging, the morphological classifications into cystic and solid types were established during early passages (P0–P2) and remained consistently stable throughout subsequent culture. Within the observation window up to the maximum achieved passage at this evaluation stage (P5), no spontaneous morphological transitions—such as cystic organoids converting into solid structures—were detected, and both phenotypes maintained their distinct structural integrity. Nevertheless, it should be noted that organoids were analyzed at early passages in this study. Prolonged in vitro culture and serial passaging may lead to clonal selection or acquisition of new mutations, potentially altering their genomic and phenotypic fidelity over time [11]. Therefore, caution is warranted when using late-passage organoids for functional or translational studies.

Our results align with prior reports in other cancers (e.g., colorectal, pancreatic, breast), where successful organoid or PDX formation often correlated with aggressive tumor biology and worse clinical outcomes [17]. However, to our knowledge, few studies have systematically evaluated such associations in HCC [19]. Notably, while successful organoid establishment itself was not predictive of survival, growth phenotype (i.e., cystic vs. solid) provided an initial indication of potential prognostic value in this cohort (Fig 5), suggesting the potential relevance of organoid morphology as a supplementary exploratory indicator of tumor behavior.

Nevertheless, several limitations should be acknowledged. First, the cohort size was modest, precluding a statistically valid multivariate analysis due to limited event counts and missing data, which prevents us from concluding morphology as an independent adverse prognostic factor at this stage. Second, although we observed a strong prognostic signal associated with organoid morphology, the biological mechanisms underlying the cystic versus solid phenotypes remain unclear. While previous studies in colorectal cancer have suggested associations with IGF1R signaling or stemness features [25], such pathways were not directly examined in our study. Third, comparative genomic analysis between cystic and solid subtypes was not feasible due to limited sample numbers and insufficient statistical power for subtype-specific variant calling. This also precluded robust investigation into whether certain mutations or driver gene alterations may preferentially associate with organoid morphology. Lastly, all organoids were analyzed at early passages, and long-term culture may introduce selective pressures or mutational drift, potentially impacting their biological fidelity [11]. Future studies with larger cohorts, multi-omics integration, and functional validation will be essential to uncover the mechanistic basis of morphological heterogeneity and its relevance to clinical behavior.

## Conclusion

Our findings suggest that patient-derived liver cancer organoids preserve the genetic and phenotypic characteristics of parental tumors and that their growth morphology, particularly the solid subtype, may reflect intrinsic tumor features and correlate with clinical outcomes within this study cohort. Given their establishment efficiency, genomic fidelity, and exploratory prognostic potential, tumor organoids represent a promising model for liver cancer research. Continued refinement of organoid culture protocols and integration with multi-omics and drug response data could enhance their translational relevance across academic, clinical, and industrial settings.

## Supporting Information

S1 Table. Somatic mutation profiles of matched primary tumor tissue and patient-derived organoid from Patient_09.

## Author Contributions

Y.S.K. and Y.H.G. contributed equally to this work, performing organoid culture, data analysis, and initial manuscript drafting. Y.S.K. and H.S.K. assisted with clinical data collection. J.S. performed histological evaluation. D.O.K. critically reviewed the manuscript, performed data verification, and contributed to manuscript revision and finalization. J.Y. and D.Y.H. supported data interpretation. W.Y. supervised organoid experiments, WES data processing and interpretation, and contributed to study design and manuscript revision. J.H.L. conceived and led the project, secured funding, and finalized the manuscript. All authors reviewed and approved the final version.

## Data Availability Statement

All relevant data are within the manuscript and its Supporting Information files. The data underlying the results presented in this study are available upon reasonable request from the corresponding authors.

