## Supplemental Table 1 for "Patient-Derived Liver Cancer Organoids Reflect Tumor Biology and Their Growth Phenotype Correlates with Clinical Outcomes"

**S1 Table. Somatic mutation profiles of matched primary tumor tissue and patient-derived organoid from Patient_09.**

Somatic variants were identified by whole exome sequencing (WES) using tumor-only variant calling with preset liver cancer gene panels. Columns indicate sample name, gene symbol, nucleotide change with transcript accession, predicted protein change, sequencing depth, number of alternative allele reads (Tumor Alt Count), and variant allele frequency (VAF). TP53 (p.Y234C) and CTNNB1 (p.K335I) were concordantly detected in both the primary tumor tissue and the matched organoid, confirming preservation of HCC driver mutations during organoid derivation. The elevated VAF of TP53 in the organoid (VAF = 1.000 vs. 0.276 in tissue) suggests clonal enrichment of the TP53-mutant population during ex vivo expansion. COL11A1 was detected exclusively in the organoid sample, possibly reflecting subclonal architecture below the detection threshold in the primary tissue or a variant acquired during culture.

| **Sample Name** | **Gene** | **DNA Change** | **Protein Change** | **Depth** | **Tumor Alt Count** | **VAF** |
| --- | --- | --- | --- | --- | --- | --- |
| **Patient_09**  **-Tis** | TP53 | NM_000546.6:c.701A>G | p.Y234C | 58 | 16 | 0.276 |
|  | LRP1B | NM_018557.3:c.5734G>A | p.A1912T | 109 | 36 | 0.330 |
|  | CTNNB1 | NM_001904.4:c.1004A>T | p.K335I | 69 | 15 | 0.217 |
| **Patient_0**  **9-Org** | COL11A1 | NM_080629.3:c.1912G>A | p.E638K | 104 | 15 | 0.144 |
|  | TP53 | NM_000546.6:c.701A>G | p.Y234C | 58 | 58 | 1.000 |
|  | LRP1B | NM_018557.3:c.5734G>A | p.A1912T | 126 | 42 | 0.333 |
|  | CTNNB1 | NM_001904.4:c.1004A>T | p.K335I | 99 | 47 | 0.475 |

*VAF, variant allele frequency; WES, whole exome sequencing; HCC, hepatocellular carcinoma.*
